# Active learning tools improve the learning outcomes, scientific attitude and critical thinking in higher education: Experiences in an online course during the COVID-19 pandemic

**DOI:** 10.1101/2020.12.22.423922

**Authors:** Izadora Volpato Rossi, Jordana Dinorá de Lima, Bruna Sabatke, Maria Alice Ferreira Nunes, Graciela Evans Ramirez, Marcel Ivan Ramirez

## Abstract

Active teaching methodologies have been placed as a hope for changing education at different levels, transiting from passive lecture-centered to student-centered learning. With the health measures of social distance, the COVID-19 pandemic forced a strong shift to remote education. With the challenge of delivering quality education through a computer screen, we validated and applied an online course model using active teaching tools for higher education. We incorporated published active-learning strategies into an online construct, with problem-based inquiry and design of inquiry research projects to serve as our core active-learning tool. The gains related to students’ science learning experiences and their attitudes towards science were assessed by applying questionnaires before, during and after the course. The course counted on the participation of 83 students, most of them (60,8%) from post-graduate students. Our results show that engagement provided by active learning methods can improve performance both in hard and soft skills. Students participation seems to be more relevant when activities require interaction of information, prediction and reasoning, such as open-ended questions and design of research projects. Therefore, our data shows that, in pandemic, active learning tools benefit students and improve their critical thinking and their motivation and positive positioning in science.

## INTRODUCTION

Academically first world countries have debated how the training of students should be, from basic primary education at schools to higher education at universities (Aldenmyr et al., 2012; Biesta, 2009; Janmaat and Piattoeva, 2007; Veugelers, 2007). A major concern is how education can collaborate in the formation of citizens and professionals capable of leading technological, economic, social, cultural and political changes (Nussbaum, 2006; Ross, 2007; White, 2013). Specifically, in the area of science, researchers should be trained with skills that go beyond the technical reproduction of experiments, but that employs critical thinking and that are capable of applying scientific concepts to propose solutions and generate knowledge (Gormally et al., 2012; Lederman et al., 2013; Turiman et al., 2012). The change of curricular programs in the STEM area (science, technology, engineering and mathematics) and new proposals for educational strategies have been stimulated in different countries (Bybee, 2010; Kennedy and Odell, 2014). Lecture-based and teacher-centered pedagogy is undergoing a shift towards more active learning, in which students build their own understanding of a subject through learning activities (Michael, 2006; Wright, 2011). The benefits of active learning seem substantial, both in cognitive learning and in the development of soft skills by students, such as leadership, problem-solving, and autonomy (Chau and Cheung, 2018; Oros, 2007; Stefanou et al., 2013). In Brazil, few efforts have been made to discuss structural changes in education from basic to university. The absence of adequate working conditions encourages teachers to adopt an old-fashioned type of education, in which passive teaching methods predominate. Although there is no state initiative that encourages the incorporation of active learning methods, some higher teaching institutions have introduced methods of problem solving, critical thinking and/ or problem-based learning with an inspiring success (Araujo and Slomski, 2013; Bestetti et al., 2014; Mesquita et al., 2015; Randi and Carvalho, 2013).

Active learning comprises approaches that focus more on developing students’ skills than transmitting information and require students to perform activities that require higher order thinking (Michael, 2006). For this, students use critical thinking, which involves analysis, reflection, evaluation, interpretation and inference to synthesize information that is obtained through reading, observation, communication or experience to answer a question (Halpern, 1999; Nelson and Crow, 2014). There are several methodologies that fit the concept of active teaching, such as inquiry-based learning, project-based learning and problem-based learning (Ishiyama, 2013; Stefanou et al., 2013; Walker, 2003). Among them is, for example, project-based learning is a model that organizes learning around projects, in which challenging questions or problems are involved that involve proposing solutions, formulating hypotheses and investigative activities (Stefanou et al., 2013).

The COVID-19 pandemic has produced a situation of health emergency, economic and social instability that challenged the entire educational system. The intense contact and exchange of information that took place during face-to-face classes in normal life have been restricted to virtual spaces. Given all these sudden changes, online courses have been a viable option to prepare students at different levels (Fig. 1). Although some groups have already reported their teaching experiences and perceptions in times of lockdown and social distance (Donitsa-Schmidt and Ramot, 2020; Dutta, 2020; Murphy, 2020; Qiang et al., 2020; Regier et al., 2020; Sunasee, 2020; van der Spoel et al., 2020), very few of them reported the impact of active learning on online courses, and rarer are the studies in post-graduate students. During the pandemic, we have seen the opportunity to validate a course model with the aim of actively encouraging students of higher education to acquire important biological concepts. We planned to create a rich, multifaceted course that integrated active learning methodologies. We incorporated active-learning strategies that allowed transit in the course from passive lecture-centered to active student-centered learning. With this approach, we were interested in understanding the added value of our course at the student formation and in answering two important questions:

**Fig. 1.**
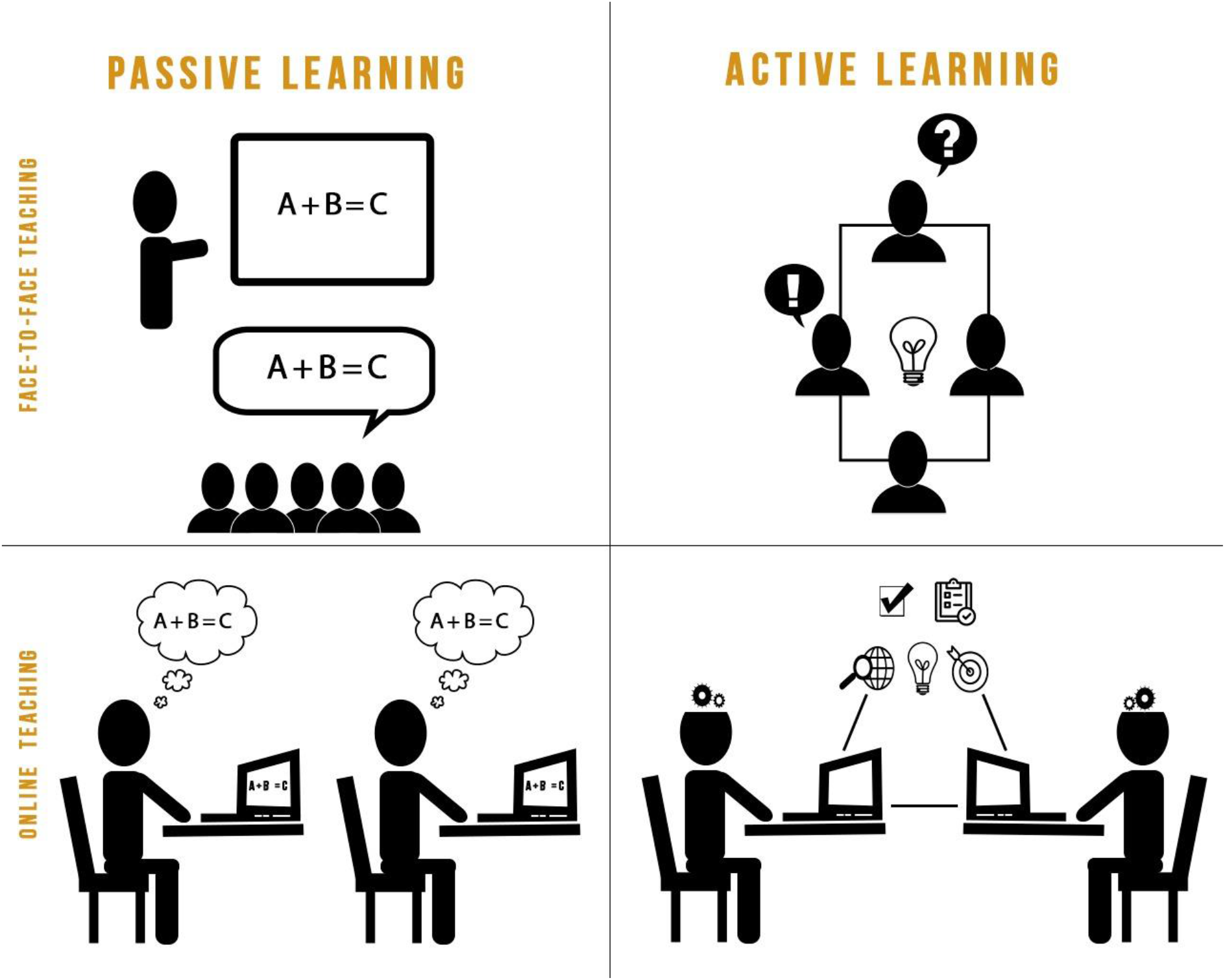
Passive (teacher-centered) and active (student-centered) learning in classroom or remote teaching models.

i. Does the course increase the cognitive and intellectual skills of the students?
ii. How was the impact of critical thinking methodologies into the student’s attitudes towards science and soft skills?

Our interest was concentrated in analyzing whether students through the course showed more enthusiasm for the concept of research and science. Crucial elements in science such as forming and testing hypotheses, defining strategies, communicating results were evaluated to determine whether critical thinking methods could improve thinking and rational logic. In order to assess students’ gains in these two aspects, we applied questionnaires to students before, during and after the course. Here, we will comment on the results of this experience that incorporated active methodologies and student-teacher interaction tools for remote higher education.

## MATERIAL AND METHODS

### 1. Undergraduate pilot course to validate online active-learning tools

In order to validate an online course model and test some active learning tools, we have offered a course aimed primarily at undergraduates. The subject of this course was cell culture which has a wide interest and application in the biological area. Knowledge on cell culture is required for some research activities and also represents a promising alternative for replacement of animal experimentation.

In order to follow contagious preventive actions during the COVID-19 pandemic, the course was administered remotely in a teleconference format through the Microsoft Teams platform. This platform allows the instructors to interact through video, audio and live chat, which gives the feeling of a personal meeting from a safe distance. Before each class, there was a moment of relaxation with “icebreaker” conversations to get to know the audience. This moment helped to create a more intimate environment and also to share tensions and concerns about the pandemic.

The course had a total of 15 hours, 10 hours of synchronous activities and 5 hours of asynchronous activities. Synchronous activities included lectures, simultaneous online quiz activities and discussion of scientific papers. Asynchronous activities consisted of two questionnaires containing guided questions for critical reading of a scientific paper (one of papers involving chronic diseases and the other infectious diseases). After returning this questionnaire, the papers were discussed during classes. To measure perceptions of the overall effectiveness of the course and the proposed methodologies, we asked students to complete a questionnaire at the end of the course.

### 2. Experimental undergraduate and postgraduate course

#### a. Course design

The experimentation course was offered as a satellite event during a symposium hosted by a Post-Graduate Program at a Brazilian state university. The focus of the course was redefined from our previous basic course to contemplate strategies for the study of infectious diseases using cell culture. In order to know the profile of the enrolled students, we applied two questionnaires containing open and closed questions: one with demographic questions and previous research experience and the other about their previous experiences with active learning methodologies.

The course had a short duration (12h total), divided between synchronous (7h) and asynchronous activities (5h). The synchronous activities of the course were structured as follows: (i) 2h of key concepts to introduce the subject and situate the content and emphasis of the course; (ii) 2h of strategies for studying the pathogen-host cell interaction using cell culture, (iii) 1h of presentation of a inquiry research project (IRP) with the subject chosen by the participant, (iv) 1h of questions about concepts and strategies to solve problems (Table 1). The “offline” time was used to prepare the scientific IRP and participate in the questionnaires with questions related to the classes. The description of the activities developed can be found in the topic “Active learning instruments/tools” below.

**Table 1.** Course Schedule.

| Learning activities | Description | S.A.T. | A.A.T. |
| --- | --- | --- | --- |
| Lecture 1. Introductory concepts | - Dynamic interaction between pathogens and host<br>- Virulence factors of pathogens<br>- Biology of the host cell<br>- Cell culture basis | 2h | 0 |
| Lecture 2. Study strategies of pathogen-host cell interaction using cell culture | - Genetic manipulation of molecules related to the interaction<br>- Changes in the host cell<br>- Secretion of factors by pathogens<br>- Evaluation of immune system modulation<br>- Evaluation of cellular communication (Protein secretion, Quorum sensing, Extracellular vesicles) | 2h | 0 |
| Simultaneous Online Quiz | Quick fixation questions answered online in real time and then corrected | 1h | 0 |
| Inquiry research project | Preparation and presentation of IRP with theme chosen by the participant | 1h | 3h |
| Problem-based inquiry | Questions of class related concepts and strategies for solving biology problems | 1h | 2h |

S.A.T.: Synchronous Activities Time. A.A.T.: Asynchronous Activities Time

#### b. Active learning instruments/tools

In order to place the student as the center of the course, we incorporated some active-learning strategies into an online course construct. Some moments of the dynamics of the classes and the approaches used during the course are gathered in Suppl. Video 1. We proposed some activities that required student’s engagement:

### i. Quiz

The quiz was a knowledge fixation tool performed at the end of lectures. In this activity, participants answered questions related to the presented content directly through the Voxvote website (https://www.voxvote.com/). Table 2 contains some examples of applied questions; the questions were corrected at the end of the time proposed by the VoxVote tool (Suppl. Video 1, min 02:36 - 02:51).

**Table 2.**
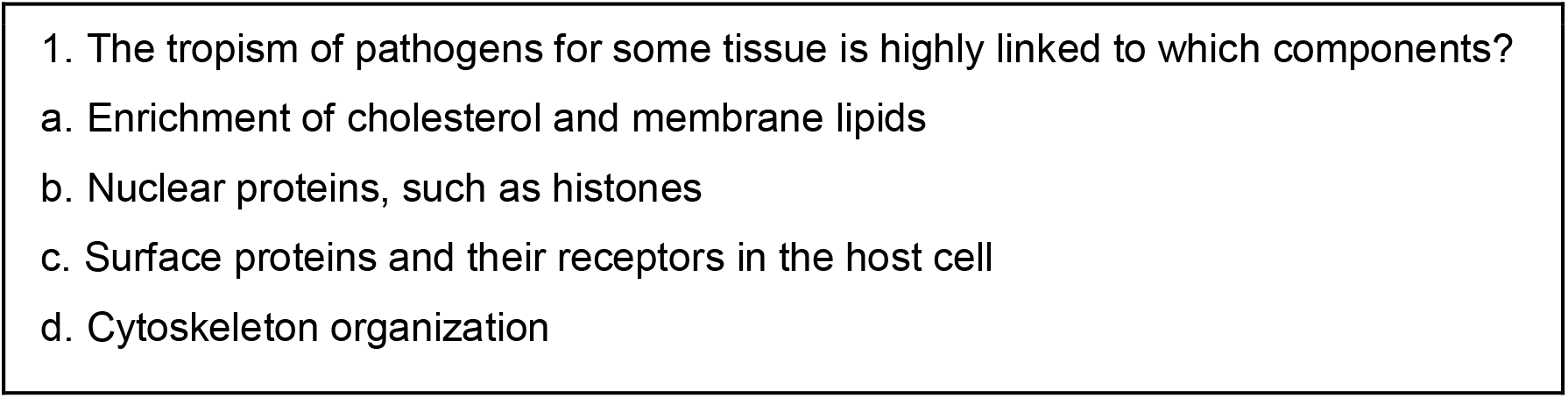
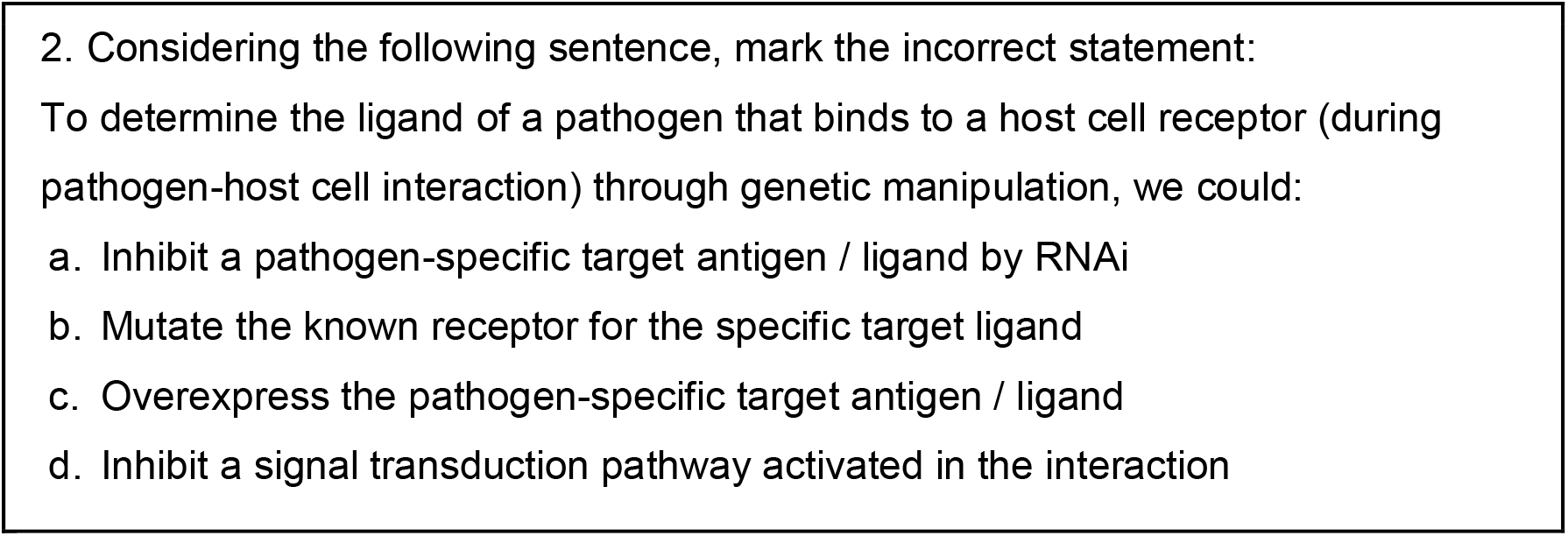
Examples of questions administered during live quiz using VoxVote.

### ii. Inquiry research project (IRP)

We proposed to the participants to develop an inquiry research project (IRP) to stimulate the construction of knowledge and critical thinking. The IRP should contain the scientific relevance of the project, main objectives and methodologies to achieve the proposed objectives. Along with the description of the project, participants could send a graphic design summarizing their project proposal, following a Graphical Abstract model indicated as a reference (Suppl. Fig. 1). The IRP was sent using Google Forms. The IRP proposals were evaluated by all instructors who selected the best 10 for presentation based on criteria of coherence and conceptualization of the biological question, ampleness of the applied methodologies and connection between the proposed strategies.

### iii. Inquiry questionnaires

Two online questionnaires were sent to all participants via email and were available for at least 48 hours. Both questionnaires contained 8 multiple-choice and 4 open-ended questions about biological concepts related to the course subject. The first questionnaire (Q1) was available before the beginning of the course, while the second (Q2) was available two days after the experimental course started. Q1 and Q2 had the same level of difficulty, with multiple-choices (basic) and problem-based questions (open-ended) (see Table 3). Q2 was answered while the students were simultaneously participating in several activities of the hosted event.

**Table 3.** Examples of questions applied in the inquiry questionnaires.

| Q1 |  |
| --- | --- |
| Multiple-choice questions | Open-ended questions |
| The fluidity of the plasma membrane increases with:<br>a) Increased unsaturated acids in the membrane<br>b) Increased saturated acids in the membrane<br>c) Increased glycolipid in the membrane<br>d) Increased phospholipid in the membrane | You read in a scientific article that Dexamethasone (a corticosteroid type drug) helps in the treatment of coronavirus infection, but the article provided only clinical data on mortality, intubation rate or length of hospitalization time. How would you investigate the mechanism of action of Dexamethasone for the benefits seen in the study? |
| Q2 |  |
| Multiple-choice questions | Open-ended questions |
| Check the correct alternative regarding the immune system:<br>a) Analyzing only one cell type can already define the complete profile of the host response during an infection.<br>b) The immune response against a pathogen is always pro-inflammatory<br>c) Dendritic cells are very homogeneous in terms of their markers and their performance.<br>d) Cytokines coordinate the action of different cell types in different tissues. Deregulation of cytokine release generates a phenomenon called cytokine storm. | One of the mechanisms of fungi and bacteria virulence is the formation of biofilm. These structures contribute to antibiotic resistance. How would you experimentally demonstrate that biofilm is important in the pathogenesis of a disease and for antibiotic resistance? |

#### c. Inquiry Questionnaire Assessment

Questionnaire responses were corrected by five evaluators. Multiple-choice questions scores were calculated by sum of the right answers. Open-ended questions required a more detailed evaluation process where four evaluation criteria were scored in each answer: comprehension, specificity, ampleness and connection. All evaluators considered whether the student had understood the question (comprehension), the approaches that the student proposed to solve the problem (ampleness), the specificity of this or these approaches (specificity) and the rationale and feasibility of the strategy (connection). For comprehension evaluation only 0 (lack of comprehension) and 1 (adequately answered). The other criteria considered three levels of score: insufficient (0), good (1) and excellent (2). The maximum score was 7 points to each answer. Answers zeroed in comprehension were not evaluated in the remaining criteria. Furthermore, the order of questions and answers were randomized to avoid possible bias during the assessment process. The scores were generated from the average of 5 evaluators. Total score was calculated by the sum of multiple-choice questions (0-50%) and open-ended questions (0-50%).

Intra-questionnaires comparisons, i.e. between questions, were assessed by ANOVA, while questionnaire differences were analyzed by unpaired t-test. All analyses were performed in GraphPad Prism version 6.01.

#### d. Analysis of students’ perception of the course

At the end of the course, students were asked to fill up their impressions and suggestions about the course in a feedback form containing multiple-choice and open-ended questions. Some questions were to choose the sentence which they felt more identified and in others the students evaluated sentences in a 5 points scale, with 0 being “nothing” and 5 being “very” (Likert scale. Likert, 1932). Open-ended questions were added to stimulate the students to express their opinion about the course. The open-ended data was coded in categories considering the most cited answer for each question. Qualitative thematic content analysis was applied to quantify answers, providing support for a quantitative evaluation.

## RESULTS

### 1. Online course can be a platform of active learning methodologies: a pilot experience

In order to validate an online course model and test some active learning tools, we have offered a course aimed primarily at undergraduates during pandemic. The wide theme Cell Culture was well received by students, attracting participants from different fields of health sciences (including biology, biomedicine, pharmacy, biotechnology and medicine - data not shown) and with different backgrounds (3,37% bachelor degree, 56.75% undergraduate students, 18.91% mastering students, 4,72% masters, 10,13% doctoral students, graduate course 2,02% and 4,05% doctoral also participated. n = 148. Suppl. Fig. 2 A).

**Fig. 2.**
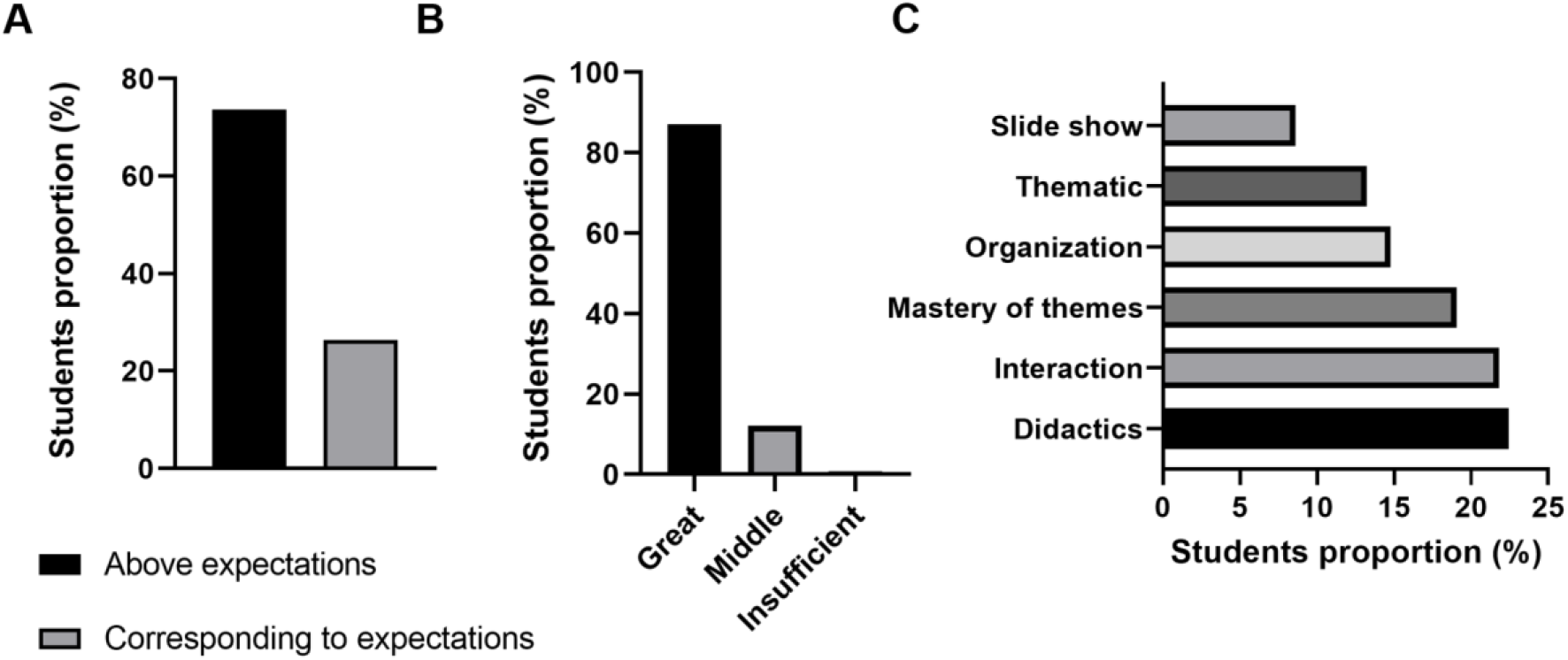
The methodology tools used during the validation course were positively evaluated by the participants. A. Course evaluation by participants. B Contribution in the course in learning cell culture. C. The open-ended question on “Course strengths” was content analyzed, and the responses were classified into categories that included similar statements.

The great advantage of remote education is being able to bring together or to mix participants from different educational institutions and different backgrounds. Participants were from 22 different Brazilian institutions and 1 foreign institution, with public and private education (Suppl. Fig. 2 B).

Although only 29% of students had worked with cell culture, the positive perception of the course was very high (Fig. 2 A). Moreover, 87% of the participants evaluated the course as excellent (Fig. 2 B). Active learning tools used during the course (real-time online quiz (live), paper reading guide, etc.) was positively rated by participants (data not shown) and the participants pointed out as main strengths the didactics, teaching methodology and the interaction between teacher and student (Fig. 2 C).

Excited by the positive experience of the first course, we decided to go deeper into a course aimed at postgraduate students to understand whether active learning tools could improve their cognitive and thinking skills. With the validation and approval of the active learning approach, we decided to maintain some activities (such as the real-time online quiz and questionnaires) and adjust some activities targeting the topic to the participants.

### 2. Experimental course: active learning tools improve the performance of students in higher education

#### 2.1 Participants Students profile: a representative sample of Brazilian higher education

The second experience with the online course model had a heterogeneous audience profile, including participants with different levels and from different locations. There were 83 enrolled, most of them master students (38.0%) (Fig. 3 A). Undergraduate students constituted 24.1% and PhD students 19.0%. There was also the participation of PhDs, constituting a very heterogeneous public. The participants belonged to 22 Brazilian institutions from different states (Fig. 3 B). Although 94% had previous research experience, only 59.4% had experience on cell culture (Suppl. Fig. 3 A), either carrying out in vitro experiments (full experience) or just accompanying other people (partial experience) (Fig. 3 C). The focus of the course was infectious diseases, which was the object of work of 59.5% of participants, including the biological model of bacteria (20.3%), fungi (15.2%), parasites (16.5%) and viruses (7.6%) (Suppl. Fig. 3 B).

**Fig. 3.**
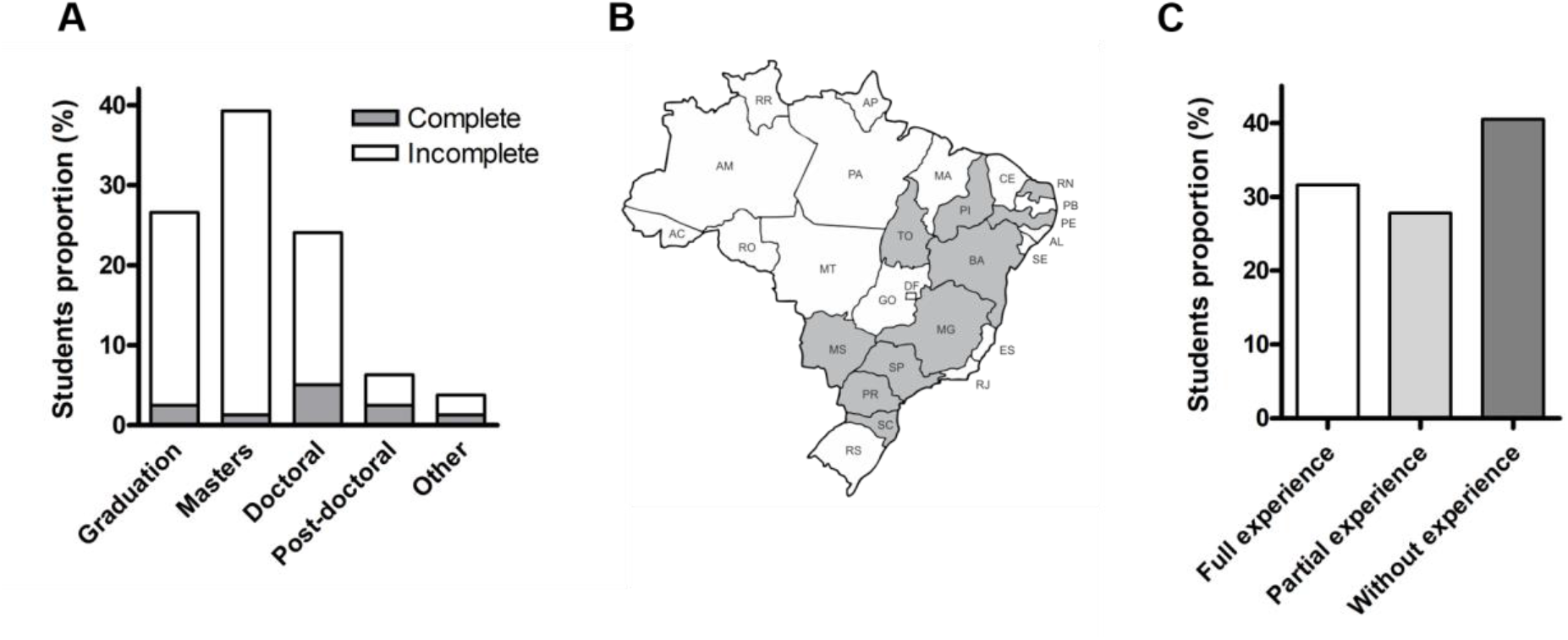
The audience of attendants to the course was heterogeneous. A. Participants’ educational background, divided between complete or incomplete. Others include “Specialist” as complete and “Incomplete second degree” and “Residency” as incomplete. B. Distribution of participants’ institutions in Brazilian states (highlighted in gray). C. Students’ previous experience with cell culture techniques.

To obtain an overview of students’ previous experience with teaching methodologies, we asked students (n=60) about which method was most used during their academic experience. The majority of the respondents (76,7%) affirm that their predominant teaching approach was passive, mainly represented by traditional lectures (Suppl. Fig. 4 A and B). Among graduate students, 50% answered that their classes have similar proportions between active and passive classes (Suppl. Fig. 4 C). When asked in an open question about what could be improved in their education (undergraduate or graduate), 72.1% of students admitted that other teaching approaches could be employed (data not shown). Most comments pointed to the necessity of interactive classes, including solving clinical cases and practical application of knowledge. The answers pointed out that most of the students (76.7%) consider that active teaching methodologies are excellent for their learning and that participating interactively in the subjects improve their apprenticeship (Suppl. Fig. 4 D).

**Fig. 4.**
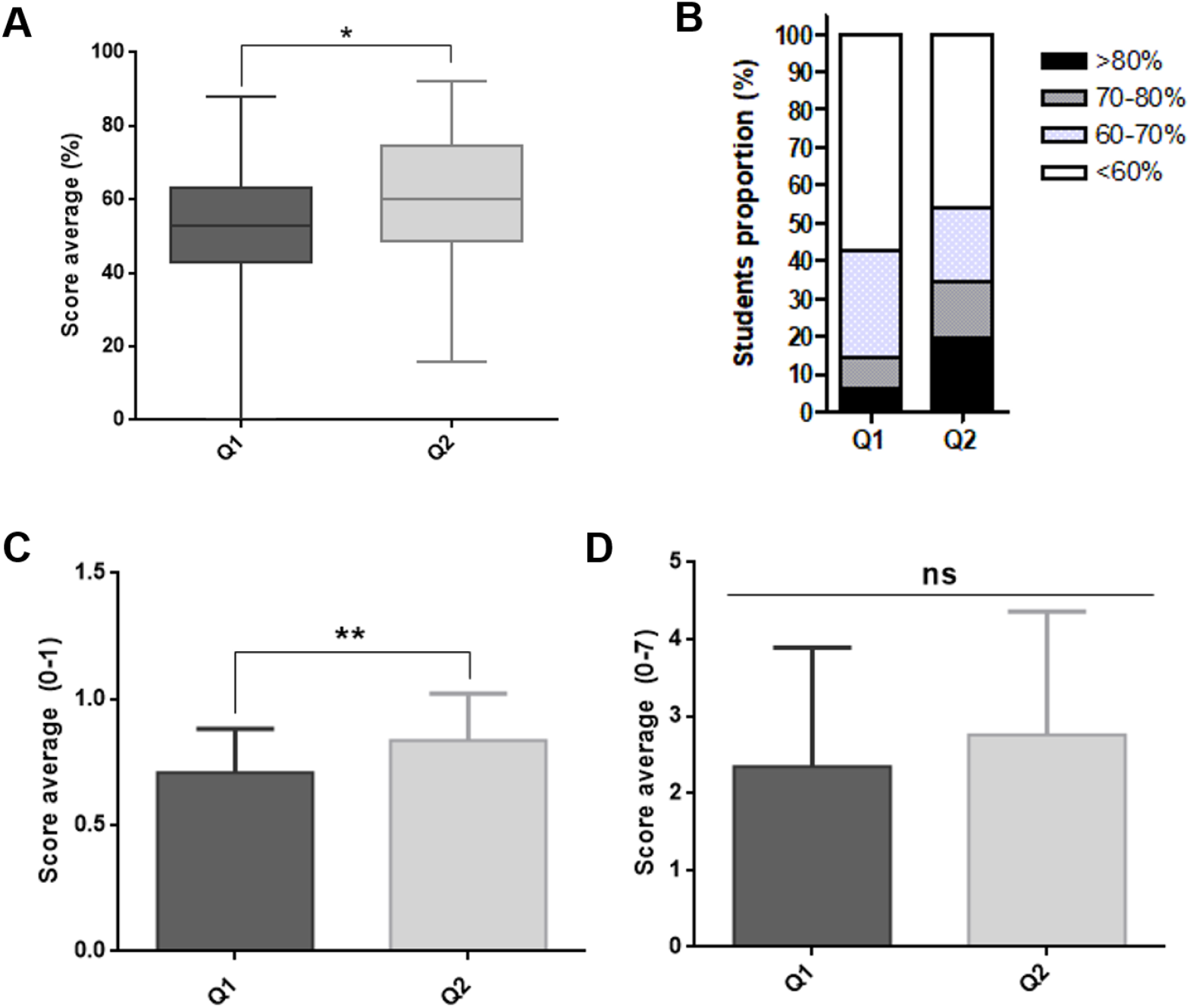
Students showed a rapid evolution in their performance during the course. A. General average score in each questionnaire (0-100%); multiple-choice and open-ended questions represent 50% of the score each; B. Proportion of students within score ranges in Q1 (n = 49) and Q2 (n=26). C. Students’ scores average only in the multiple-choice questions between questionnaires; D. Students’ scores average only in the open-ended questions between questionnaires.

#### 2.2. Active methodologies promote improved short-term learning outcomes

Interested in observing the development of students during the course, we used research-based learning approaches through the application of questionnaires in a pre-test (Q1) and post-test (Q2). Fifty-four students participated in the online questionnaire activities (Suppl. Fig. 5 - graduation: n=14; masters: n=25; doctoral: n=14; post-doctoral: n=3; other: n=1). Most of them participated in the first questionnaire (Q1: n=49), while a minority participated in the second one (Q2: n=26); finally, twenty students participated in both questionnaires.

**Fig. 5.**
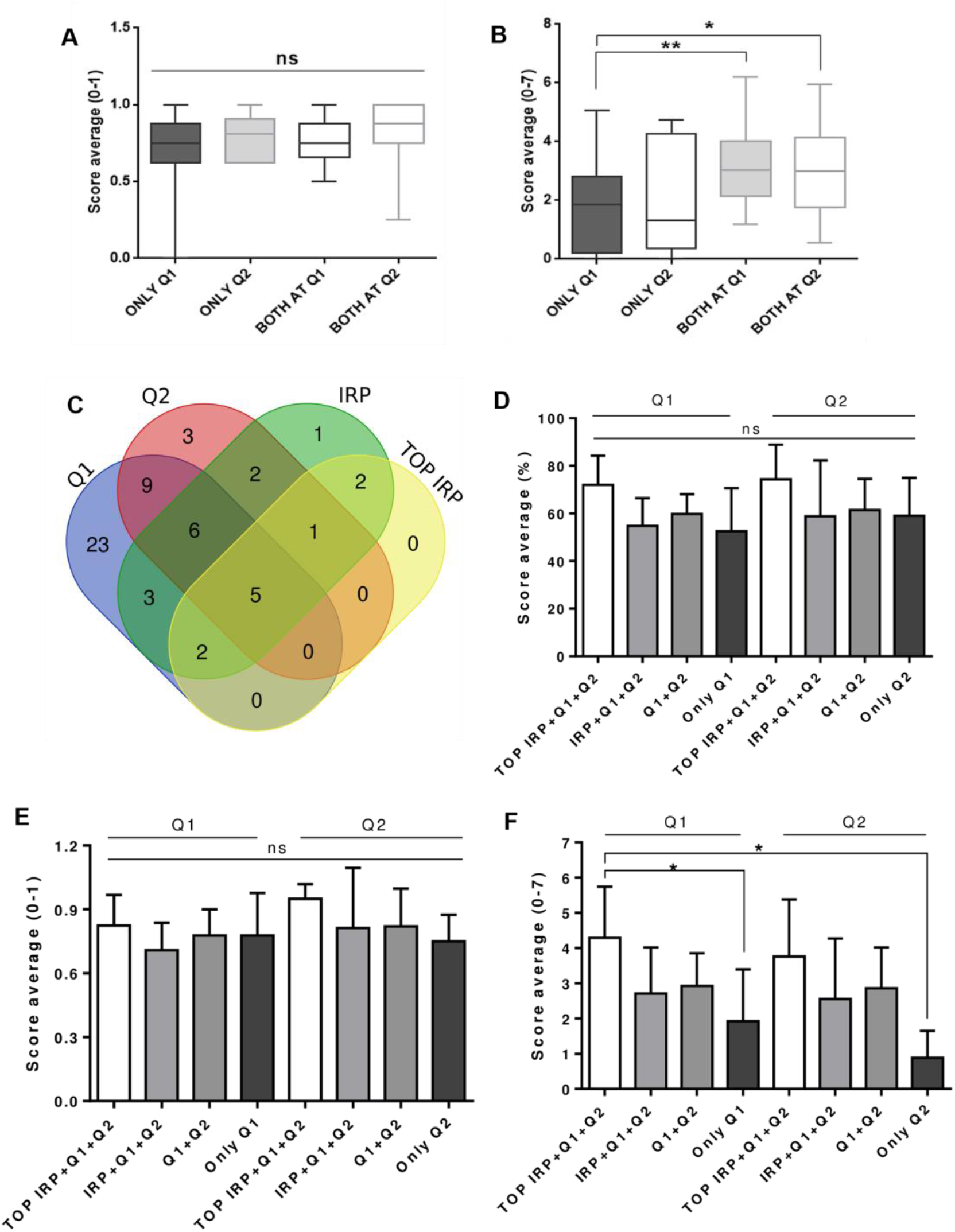
Highly engaged students have better performances in open-ended questions. A. Students’ scores average in the multiple-choice questions within each engagement subgroup; B. Students’ scores in the open-ended questions within each engagement subgroup. C. Venn Diagram, representing the number of participants in each activity (Q1, Q2, IRP and TOP IRP). D. Total score (%) in Q1 and Q2 analyzed in groups classified by the level of engagement in the course activities. The questionnaire to which the average scores refer is indicated by the horizontal bars (Q1 or Q2). E. Students average in multiple-choice questions within each group engagement. F. Students average in open-ended questions within each group engagement.

The average scores of students in the questionnaires was higher in Q2 compared to Q1 (Fig. 4 A, Suppl. Table 1). This progress was distributed similarly through multiple-choice (13,51%) and open-ended questions (15.67%). In average, no student had zeroed their score in Q2, which may represent that students were more committed to the second test (Fig. 4 A). The proportion of students with high performance (total score > 80%) was at least 3 times higher in Q2 compared to Q1 (6,1% at Q1 and 19,2% at Q2. Fig. 4 B). The students showed improvement in all four criteria evaluated in the open-ended questions from Q1 to Q2 (Suppl. Fig. 6). When multiple-choice and open-ended questions were analyzed separately, Q2’s superior performance was predominantly due to the scores at the multiple-choice questions (Fig. 4 C) than from the open-ended (Fig. 4 D).

**Fig. 6.**
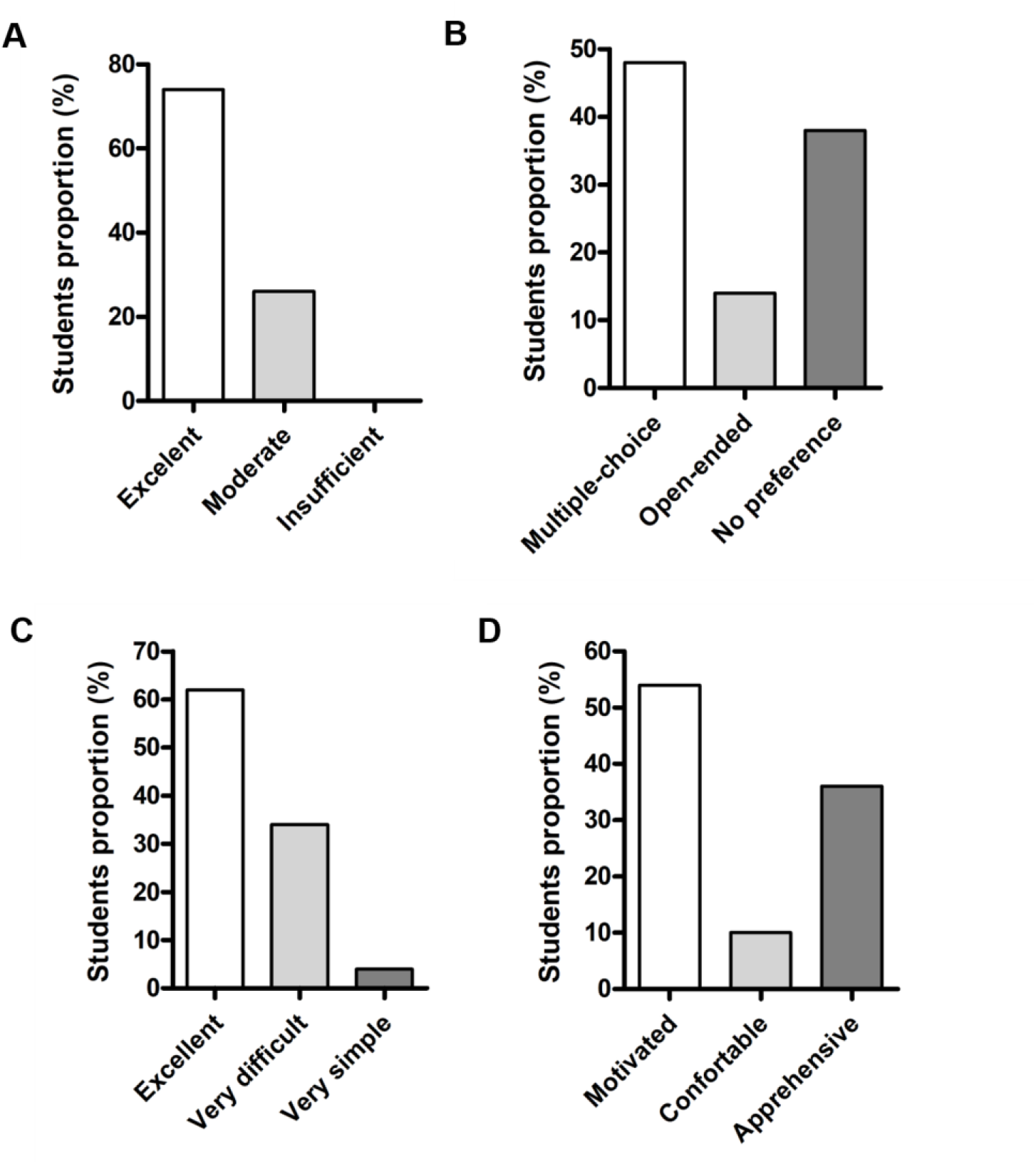
Students demonstrate a positive feeling about active learning tool. A. Percentage of responses from students on the multiple-choice question “How do you think the course contributed to your learning?”, With possible answers “Excellent”, “Moderate”, “Insufficient”. B. Percentage of responses to the multiple-choice question “In the questionnaires, what type of question do you prefer?”, With possible answers “Multiple- choice”, “Open-ended”, and “I have no preference”. C. Percentage of students’ responses to the question “How do you evaluate the problem-based questions present in the questionnaires?”, With possible answers “They were excellent”, “They were very difficult” and “They were very simple”. D. Percentage of responses to the question “How did you feel during the conduct of the inquiry research project?”, With possible responses being “Motivated”, “Comfortable”, and “Apprehensive”. The percentage of responses was calculated on the number of students who answered the questionnaire (n = 50).

We hypothesized that the overall improvement of open-ended questions may be due to a lower engagement at the more difficult and exploratory questions (such as the open-ended questions). Regarding this point we calculated the student dropout rate to each question by the rate of NA answers - i.e. described as blank answers and “I don’t know” type of answers. In fact, the dropout for open-ended questions (Q1: 22% and Q2: 15%, Table 4) was higher than for multiple-choice questions, which was irrelevant (Q1: 2% and Q2: 0%). Furthermore, there was a 31.8% reduction in the dropout rate in open-ended questions from Q2 compared to Q1 (Table 4). This may indicate that the students felt more confident and motivated to commit intellectual effort during the performance of Q2, resulting in a better outcome.

**Table 4.** Dropout rate among open-ended questions in Q1 and Q2. Dropout was considered for blank answers and “I don’t know” type of answers. “OE” stands for open-ended questions from 1 to 4 in each questionnaire.

|  | OE 1 | OE 2 | OE 3 | OE 4 | Dropout average |
| --- | --- | --- | --- | --- | --- |
| Q1 | 14% | 32% | 22% | 20% | 22% |
| Q2 | 19% | 11% | 7% | 22% | 15% |

**Table 6.**
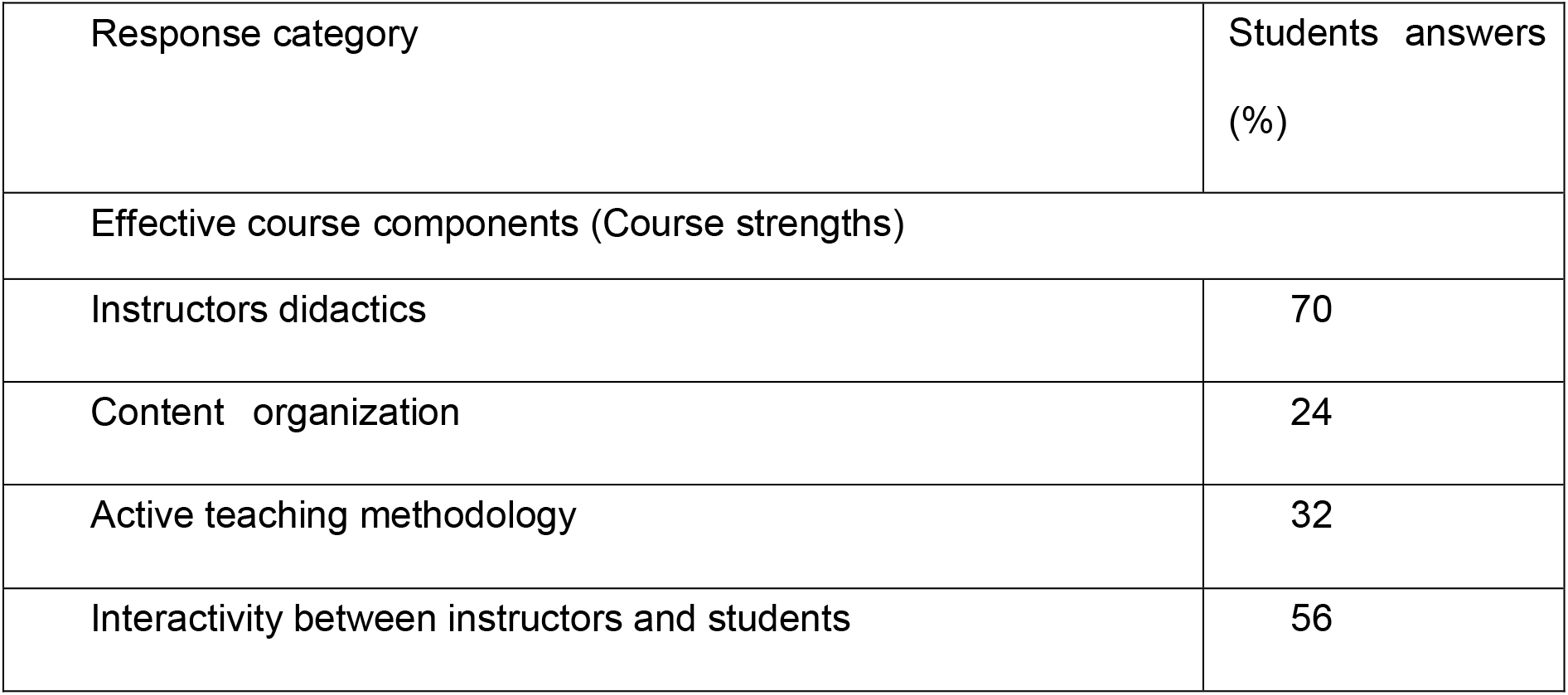
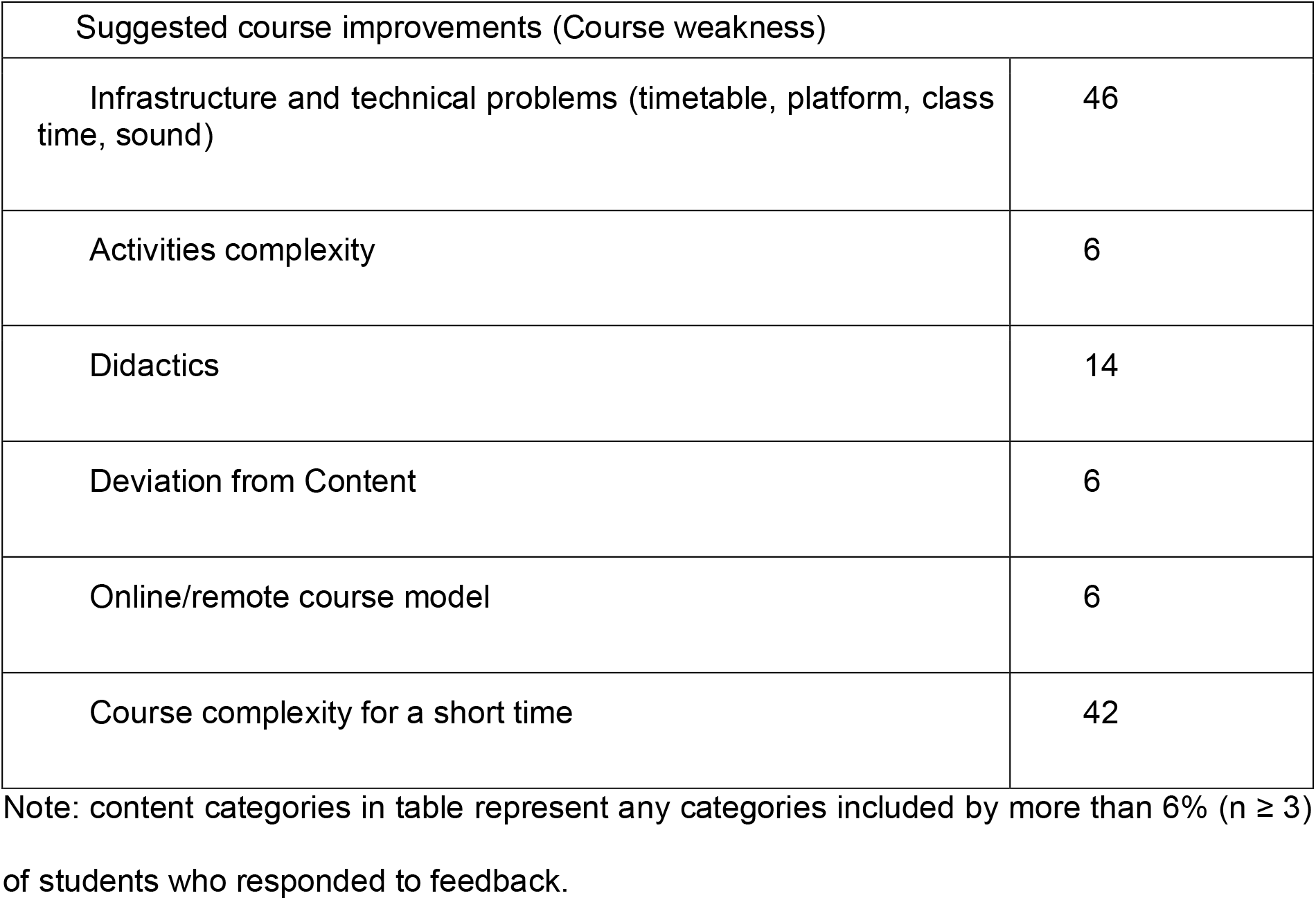
The main positive points cited by the students were didactics, teaching methodology and instructor-student interaction. The open-ended questions on “Course strengths” and “Course weaknesses were content analyzed, and the responses were classified into categories that included similar statements.

#### 2.3. Formulating hypotheses and proposing strategies: a scientist-like experience through project-based learning

The inclusion of project-based learning strategies is effective in STEM courses, to involve students in authentic “real world” tasks (Reeve and Tseng, 2011; Stefanou et al., 2013). During the course, students were motivated to prepare a mini scientific project to answer a biological question of their interest, applying cell culture strategies (see Materials and Methods). The elaboration of the scientific IRP represented the most demanding activity for the student and we had only 26,5% of participation (22 IRPs), most of which are master’s students (Suppl. Fig. 7). This type of activity was a challenge for the students, who feel freedom to “think outside the box” and find ways to answer their biological questions. Many students elaborate different and curious hypotheses, from which the instructors selected the 10 best IRPs based on criteria of coherence, conceptualization, applied methodologies and connection between the proposed strategies. We were able to see some students who stood out for the quality of their IRP proposal. Interestingly, among the 10 best IRPs selected, the fourth part was written by undergraduate students (data not shown). In addition, we reserved a period of the course for presentation of the selected IRPs to the whole class at a “symposium-like moment”, using their graphical abstracts as a support. This type of activity adds other soft skills to students, such as communication and accepting challenges, essential for future scientists. Part of the presentations of the selected students and their graphical abstract / poster as other course activities were compiled in Suppl. Video 1 (min 03:01 - 03:51).

**Fig. 7.**
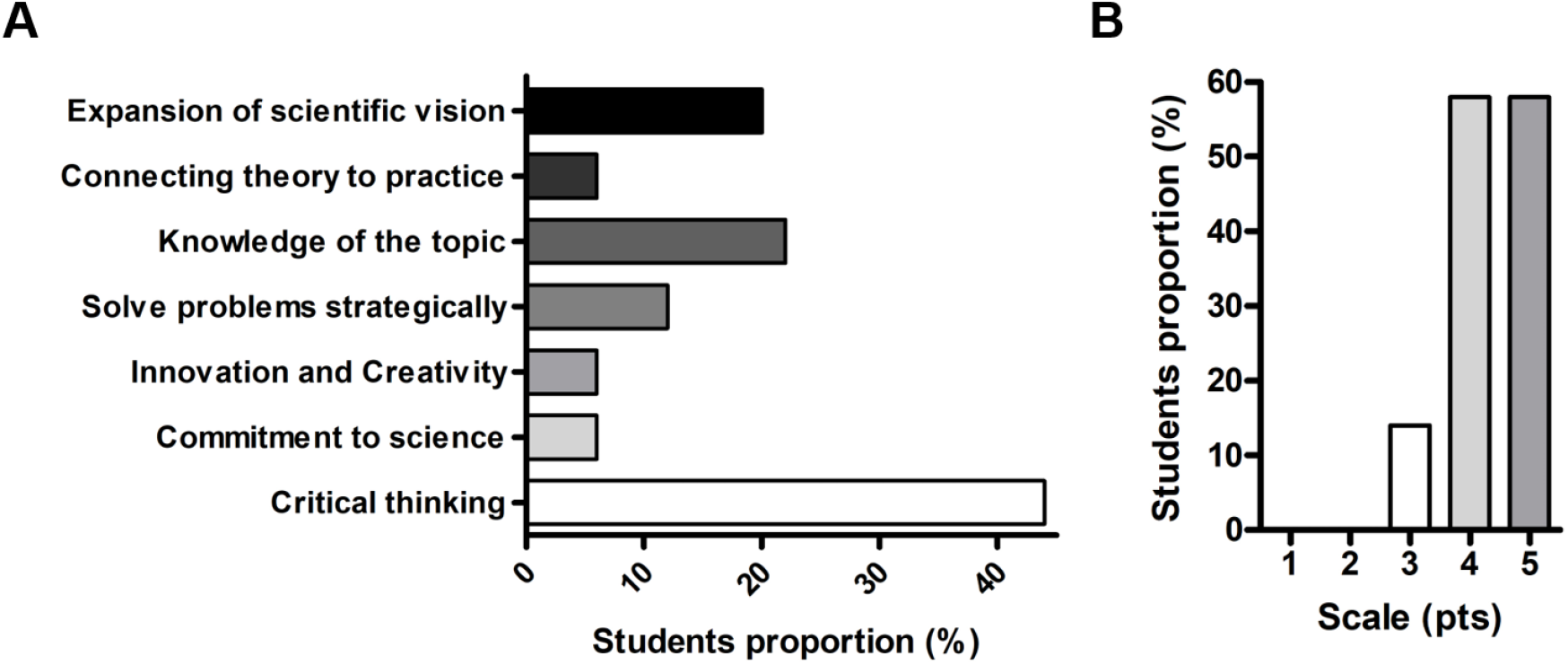
Active methodologies are able to increase the incorporation of knowledge, motivation in front of science and students show gains in soft skills. A. Answers to the open question “what are the main gains you obtained with the course?” were categorized among common themes (showing categories that comprise 6% (n = 3) or more of the answers). B. Student responses to the question “how motivated are you to solve scientific problems using critical thinking after the course?” on a scale of 1 to 5 (1: nothing; 5: very). The percentage of responses was calculated on the number of students who answered the questionnaire (n = 50).

#### 2.4. Engagement in active-learning activities correlates to better student performance

Active methodologies place the student as the center of learning and for this reason their effectiveness relies heavily on the student’s engagement in activities. Motivated by the various studies that show a positive correlation between student engagement and performance (Kennedy and Odell, 2014; Plump and LaRosa, 2017; Renninger and Hidi, 2006; Rodenbusch et al., 2016; Strayhorn, 2008; Wiggins et al., 2017), we assessed whether the most engaged students during our course had higher scores.

First, we evaluated the scores of the group of students who participated in both inquiry questionnaires (“BOTH”) separated from those who have answered only one of the questionnaires (“ONLY Q1” or “ONLY Q2”). This analysis showed that students that were engaged in both activities had higher performance in open-ended questions, but not in multiple-choice (Fig. 5 A, B).

We hypothesized whether engagement in activities proposed during the course (questionnaires and IRP) would be related to the best performance of students. For this, we considered the following groups: students who had been selected as TOP IRP and also participated in both Q1 and Q2 (TOP IRP + Q1 + Q2, n=5), students who participated in Q1 and Q2 and sent IRP (but were not TOP IRP, named IRP + Q1 + Q2, n=6), students who only participated in the questionnaires (BOTH Q1 and Q2, n=9) and those who participated in only one of the questionnaires (Only Q1, n=23 or Only Q2, n=3) (Fig. 5 C). Interestingly, half of the students who were selected as TOP IRP also engaged in both Q1 and Q2 (n = 5).

The students who participated in all activities had higher score levels when compared to the other groups of engagement, mainly in the open-ended questions analyzed separately (Fig. 5 D). Among the students who participated in the IRP, the best scores were from the students who were in the top-ten IRP (TOP IRP) (Fig. 5 C and D). Our data show that the participants who answered only one of the questionnaires (Only Q1 or Only Q2) had the worst scores in the open questions and shows that involvement in more than one activity improves the student’s performance (Fig. 5 D). Altogether, the data show a positive trend in the relationship between engagement and performance (Suppl. Fig. 8).

#### 2.4. Active learning tools improve students’ critical thinking and motivation in science

The evaluation of the course was positive by 74% of the participants (n=50), who considered that the course was excellent (Fig. 6 A). The open-ended questions on “Course strengths” and “Course weaknesses were content analyzed, and the responses were classified into categories that included similar statements (Table 6). Among the strengths, 70% of the students considered the didactics as a strong point, which includes the quality of the presentations, the confidence of the instructors regarding the domain of the content, the lesson plan, and the dynamics of the class. 56% of the participants assessed that the student-teacher interaction was a positive aspect of the classes, where the students revealed that they felt included (even remotely). Another point highlighted as strengths of the classes was the teaching methodology and the subjects covered, which brought a balance between variety and depth. As negative points of the course, issues with infrastructure and technical problems (such as timetable, platform, class time, sound) and course complexity for a short time were mentioned.

The students’ feelings about the course’s active learning tools were assessed by the feedback form. Students were instructed to rate from 1 to 5 on how positive the online inquiry questionnaires were for their learning, being 1 “negative” and 5 “very positive”. The average score of the responses was 4.34, indicating that the questionnaires were validated by the students. Regarding the type of question contained in the questionnaires, 48% of students prefer multiple-choice questions (Fig. 6 B). This shows that at least half of students prefer questions that students prefer questions that only recall information and do not require elaborating their own reasoning. Despite the high preference for multiple-choice questions among the participants (48%), 62% considered that discursive problem-solving questions are a great way to make them think critically and formulate strategies for real situations that a researcher faces (Fig. 6 C).

One of the proposed activities was the writing of an IRP about some biological question of their interest. The participation rate in IRPs was relatively low (26.2%), being 68,2% postgraduate students. Interestingly, 54% of the participants felt motivated during elaboration of the scientific project (Fig. 6 D). In an open question, the participants affirm that the elaboration of an IRP improves its positioning in science, becoming more critical and more motivated. It is also mentioned that the IRP stimulates the acquisition of more knowledge, they are able to expand their scientific vision, simulate a real situation of researchers and collaboration in scientific communication (data not shown).

In general, there was a demonstration of positive perception regarding active learning methodologies by most students (96%) (data not shown). The main points commented by the students regarding their perception of active methodologies were that they are more effective for lasting learning, stimulate critical thinking and improve the dynamics of the class and the student-teacher interaction.

When consulted in an open-ended question about the skills they improved with the course, the answers were directed to three points: incorporation of knowledge, motivation about science and gains in their skills on scientific processes. 44% of the participants cited an improvement in their logical critical and rational thinking. A gain in knowledge of the subject was pointed out by 22% of them, and the expansion of the vision by 20% (Fig. 7 A). It is also interesting to note that 94% of the participants indicate that the course was able to give a real insight into problems that scientists face in their research. When questioned how motivated they are to solve scientific problems using critical thinking after the course on a scale of 1 to 5 (1: nothing; 5: very), the average response was 4.14, with a rating of 4 and 5 by 86% of them (Fig. 7 B).

## DISCUSSION

The constant concern with excellence in the scientific training of academics encountered a new challenge during the COVID-19 pandemic: how to engage students in effective learning in remote education? This question was the driving force of our study, which reports a semi experimental online course for higher education. Our course incorporated active methodology tools that promoted the integration of students in the construction of knowledge and stimulated their critical thinking skills. For this, we proposed problem-based learning strategies in questionnaires, elaboration of a scientific project and online quiz in order to complete the lectures. In the last few months, there has been a huge increase in the number of studies dedicated to developing and validating active learning strategies in remote or hybrid education, driven by the pandemic (Ghazi-Saidi et al., 2020; Hasnine and Ahmed, 2020; McGreevy and Church, 2020; Murugesan and Chidambaram, 2020; Qiang et al., 2020).

Our study was interested in evaluating mainly two types of achievement in students: i. Cognitive and intellectual skills (learning outcomes) and ii. Critical thinking, attitudes towards science and soft skills. For this, different activities and questionnaires were applied before, during and after the course. Our data show that student engagement in the different active learning tools proposed is directly linked to their performance in the course. The average score of the groups that participated in all the proposed activities and stood out in the writing of the IRP was considerably higher compared to the groups with less involvement in the course when evaluating the discursive questions. In fact, other studies have already shown that active learning approaches in the classroom improve academic performance. In a long-term study (3 years), the implementation of problem-based learning (PBL) and learning by teaching (LbT) resulted in an increase from 5 to 6-7 in the average scores in final exams of engineering students (Freitas et al., 2016). Interactive-engagement also shows score improvements in physics courses compared to traditional pedagogical strategies (Hake, 1998).

Our data show that student involvement is a key point for their learning. This is widely accepted and experienced at different levels (Prince, 2004; Reeve, 2013). Emotional, behavioral and cognitive dimensions can be considered when analyzing engagement (Parsons et al., 2014). First, emotional engagement happens when students are emotionally affected and motivated by the learning environment (Blumenfeld et al., 2012). In our courses, introductory icebreakers and friendly communication was a factor that contributed to students to feel comfortable in interacting with instructors and with each other. Second, behavioral engagement corresponds to attitudes students demonstrate in class, such as listening and paying attention to the class or the persistence and concentration in activities (Fredricks et al., 2004). In this scenario, at least three forms of interaction were provided (chat, audio only and video), in which the chat demonstrated that students were constantly connected to instructors during the presentation. Finally, cognitive engagement happens when students apply their ability to select, connect and plan in constructing and self - regulating the learning process (Corno and Mandinach, 1983; Parsons et al., 2014). Here, these movements were detected, under our point of view, in the construction of the inquiry research project and in the responses to open-ended questions in both questionnaires, in which students provide strategies to real problems inside and outside their fields of study. All three dimensions of engagement are linked together and may contribute to improvement on students’ academic performance, then one should not consider them solely.

Beyond the intellectual benefit, traditionally used as teaching quality indicators, we hypothesized that student-centered teaching methodologies would lead to a positive attitude or perception with science and thinking skills. In a self-assessment, students reported that they had an improvement in their critical thinking, which involves judging the information with criteria and healthy skepticism. This relationship between active learning and improving critical thinking has been reported in other groups around the world (Kim et al., 2013; Nelson and Crow, 2014; Styers et al., 2018). Active-learning strategies (such as collaborative work in small groups and case studies) improved students critical thinking skills as measured by the Watson-Glaser Critical Thinking Appraisal, which assesses decision making ability as well as predicts judgment, problem solving, and creativity (El Hassan, 2016). Umbach and Wawrzynski (2005) analyzed two sets of American national data and showed a positive relationship between university environments where teachers used active and collaborative learning techniques and students’ gains in personal-social development. Improving students’ ability to recognize problems and apply effective strategies and solutions to fundamental challenges in the field is the basis of good scientific training. Our results show that tools of active methodology can impact the attitude of students that will be reflected in future scientists able to position themselves in the face of problems.

The improvement of the indicators added to the approval of the course by the students confirmed that the approaches were well chosen and encouraged us to write our experience in order to facilitate the implementation of active methodologies in other courses. We opted for active learning tools that could be easily applied to the virtual environment, improving the dynamics of the classes. Online questionnaires seem to be a great option for validating students’ learning, and makes them reflect on the class and apply their knowledge in the answers. Because our courses aim at a scientific formation associated with the resolution of real problems, the questionnaires addressed both concept questions and interpretive/exploratory open-ended questions. This allowed us to highlight a clear problem in Brazilian education: students are trained as “information recorders / archivers” and not as “critical thinkers”, as many students showed good levels in concept questions and poor performance in problem-based questions. The use of open and closed questions is ideal to provide greater freedom of responses for students and to stimulate reasoning, but they also need clear criteria for their correction. In order to guarantee the impartiality of the corrections, all five instructors of the courses corrected all the questions and the scores were given by an average between evaluators.

During the course design, we were interested in getting immediate feedback on student learning in relation to the main concepts discussed. For this, at the final of everyday classes, approximately 10 final minutes were reserved for an online quiz. This activity was very interesting to reaffirm “take home messages”, that is, what the student cannot “get out of class” without learning and their perception about the acquired knowledge. There are several online tools for this type of quiz, and we emphasize that the most interesting ones are those that allow a real-time assessment of the result with a percentage of “votes’’ in each of the questions. This allows questions to be promptly corrected and students can use that time to clear up any doubts.

During the undergraduate course, we opted for a questionnaire that represented a “critical reading guide” for scientific papers. Participation in the questionnaires was very positive, but we replaced this activity with the elaboration of a mini-scientific project in the graduate course, since reading scientific papers is a basic/trivial activity for graduate students. The preparation of the IRP represented the most demanding activity for the student, because there he should use the knowledge of the course to answer a scientific question of his preference. This type of activity gives students freedom to “think outside the box” and search for ways to answer biological questions that interest them. With this activity, we detected - observed some students who were highlighted for their commitment to develop a project as a principal investigator. In addition, we reserved a time within the course for some students who had the IRPs selected to present for everybody. This type of activity adds other soft skills to students, such as communication and accepting challenges, which are essential for future scientists.

Although we have achieved good results as an online course model for higher education, we have encountered some limitations in our study. The course was presented in a short time (3 inconsecutive days) which hampered a robust evaluation regarding the impact of active tools in student progress. Also, the experimental course was transmitted simultaneously with other activities of the hosted congress, which may have impacted on students’ outcomes due to other demanding activities. In addition, because it is an optional course (as a satellite event), there were no ways to require student participation, nor condition performance to the approval of the course. This could have been caused, among other possible reasons, by the low responsiveness in certain activities, showing that part of the students only engages in activities when they are requirements for approval. Previous experiences with the theme were not considered as a differential advantage, students from different fields in health and biological sciences were analyzed together; the same happened to undergraduate students and postdoctoral fellows, for example. Finally, a point that can be seen in a positive and negative way was the heterogeneity of the class. This was interesting because it brought the most different backgrounds to the same class, however, it also made it difficult to know about the level of knowledge among students, since the same knowledge could be very basic or essential for some and very advanced or specific for others.

Interestingly, although our study was carried out during a pandemic, with a limited number of students, our data reflect the profile of Brazilian education. The students admit that most of their academic training was with passive approaches, but they are interested and willing to more interactive activities. This exposes a gap in the unequal Brazilian educational model: changes in the educational environment are strongly necessary to prepare citizens socially and personally able to participate in society in a democratic way (UNESCO, 2004, McCowan, 2007, INEP 2003). The current model of higher education in force in Europe after the establishment of the parameters determined by the European Higher Education Area prioritizes among the student’s abilities the development of an autonomous learning capacity (Boni and Lozano, 2007; González, 2008). However, the models found in traditional schools, including Brazil, prepare students equally, minimizing the idea that knowledge acquisition is motivated in cognitive, personal and also social skills (Morán, 2015).

The introduction of active learning methodologies has been widely encouraged worldwide, but it requires a great effort from both teachers and students: educators need to review their lesson plans and add new tools and students need to be willing to engage in the construction of knowledge (Wright, 2011). Unquestionably, the process requires dynamic instructors, with a flexible mind and willing to use the class to produce a transformation in the students to acquire knowledge through active methodologies. The course was carried out after three months in full lockdown. We have no elements to evaluate if the impact of our proposal could be different spending more time within the course. Certainly, with all the uncertainties of this moment in the world, our experience reaffirms the remote method of learning when using elements of critical thinking and active methodologies, with a real benefit of self-confidence and empowerment of students to motivate themselves in their long and arduous road to be a scientist.

Above all, our experience showed that making the student the center of the class brings not only cognitive benefits (such as intellectual growth) but also in the psychosocial and personal spheres, giving students independence, improvement in their effective communication and in their ability to accept challenges for self-development. Our data shows that active learning tools that require constant engagement benefit students and improve their critical thinking. This study also shows that if courses on various scientific topics were reformulated by adding active methodologies, it is likely that more students will obtain better intellectual baggage and positive positioning towards participation in science, forming/preparing more powerful thinkers.

## Acknowledgments

We are grateful for the invitation from the Graduate Program in Biosciences and Pathophysiology (State University of Maringá) through Prof. Gessilda de Alcantara Nogueira de Melo to participate in the VII International Meeting of Biosciences and This study received support from FIOCRUZ, UFPR, CNPq, CAPES M.I.R is currently fellow from CNPq-Brazil

