## Supplementary material for "Active learning tools improve the learning outcomes, scientific attitude and critical thinking in higher education: Experiences in an online course during the COVID-19 pandemic": Table and figures

**SUPPLEMENTAL MATERIAL**

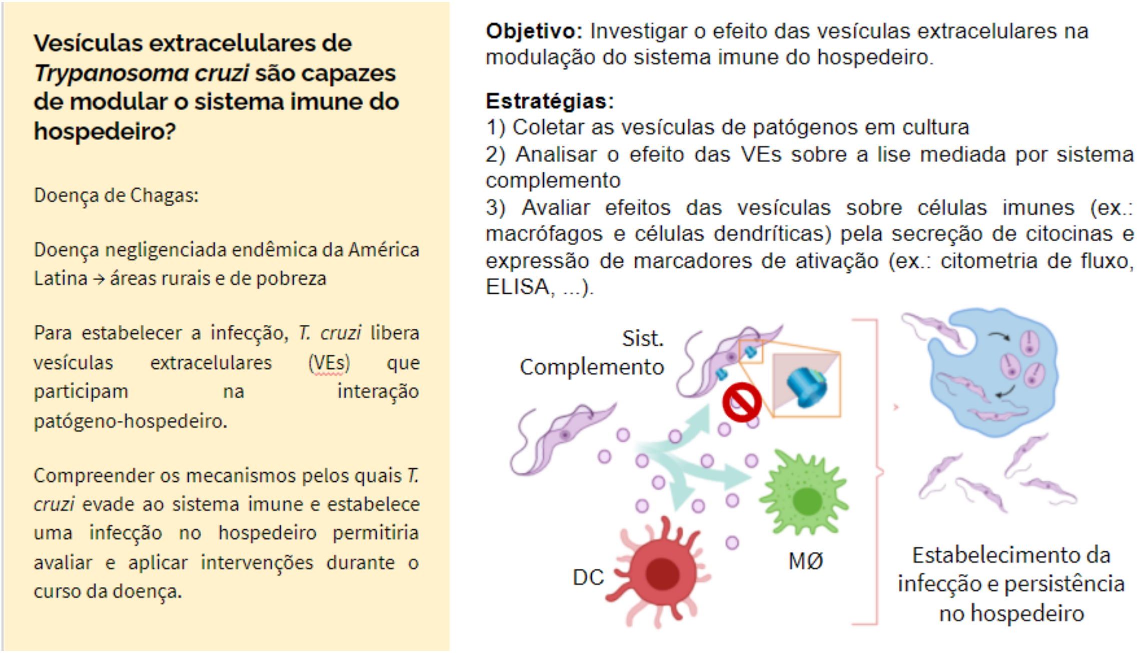


**Suppl. Fig. 1. Example of graphical element.** Initially, students incorporate the theme and importance in public health. Describe the objectives and relevance of the research, in addition to the proportions to solve the problem addressed.

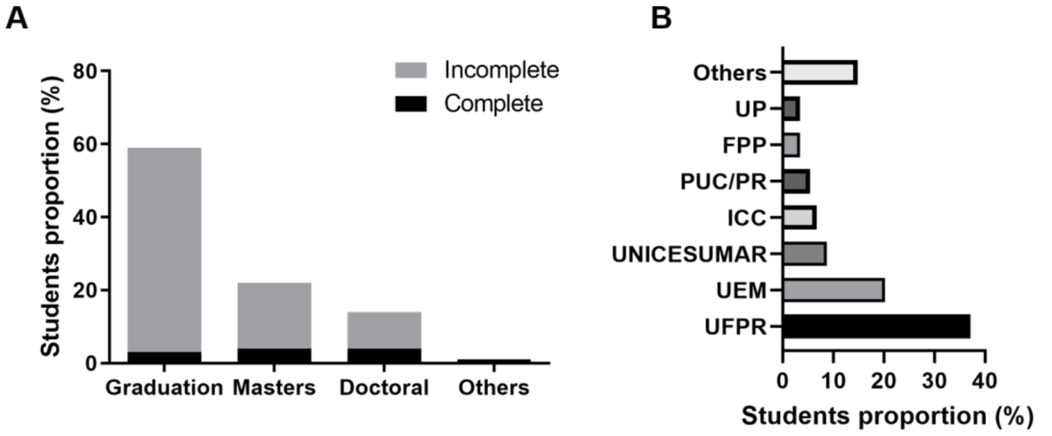


**Suppl. Fig. 2. Characterization of participants in the cell culture course.** A. Education level of participating students. B. Education Institute of participants. Federal University of Paraná (UFPR), State University of Maringá (UEM), Carlos Chagas Institute (ICC), Pontifical Catholic University of Paraná (PUC/PR), Pequeno Príncipe College (FPP) and Positivo University (UP).

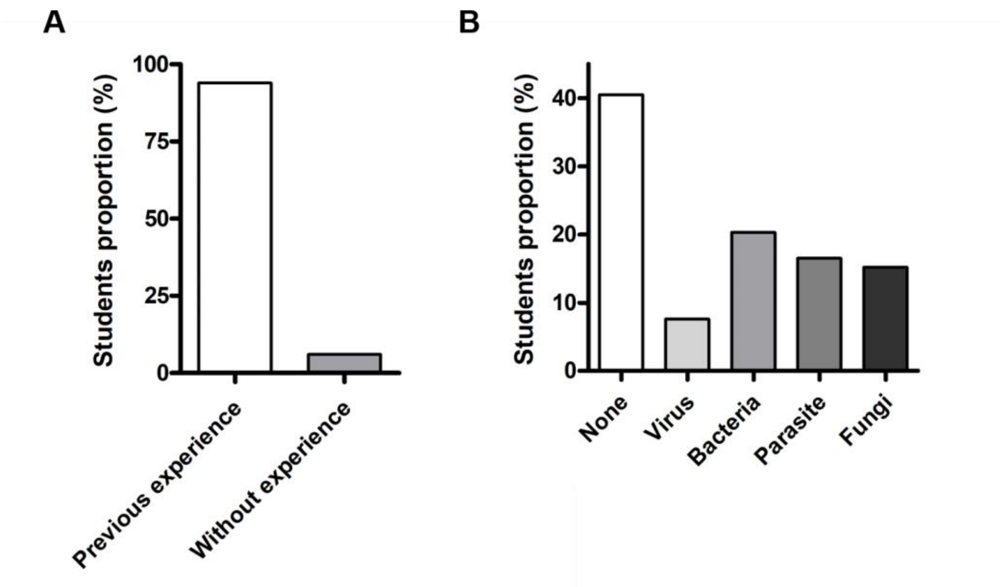


**Suppl. Fig. 3. Characterization of participants in the experimental course.** A. Participants previous experiences with research. B. Biological model investigated by course participants.

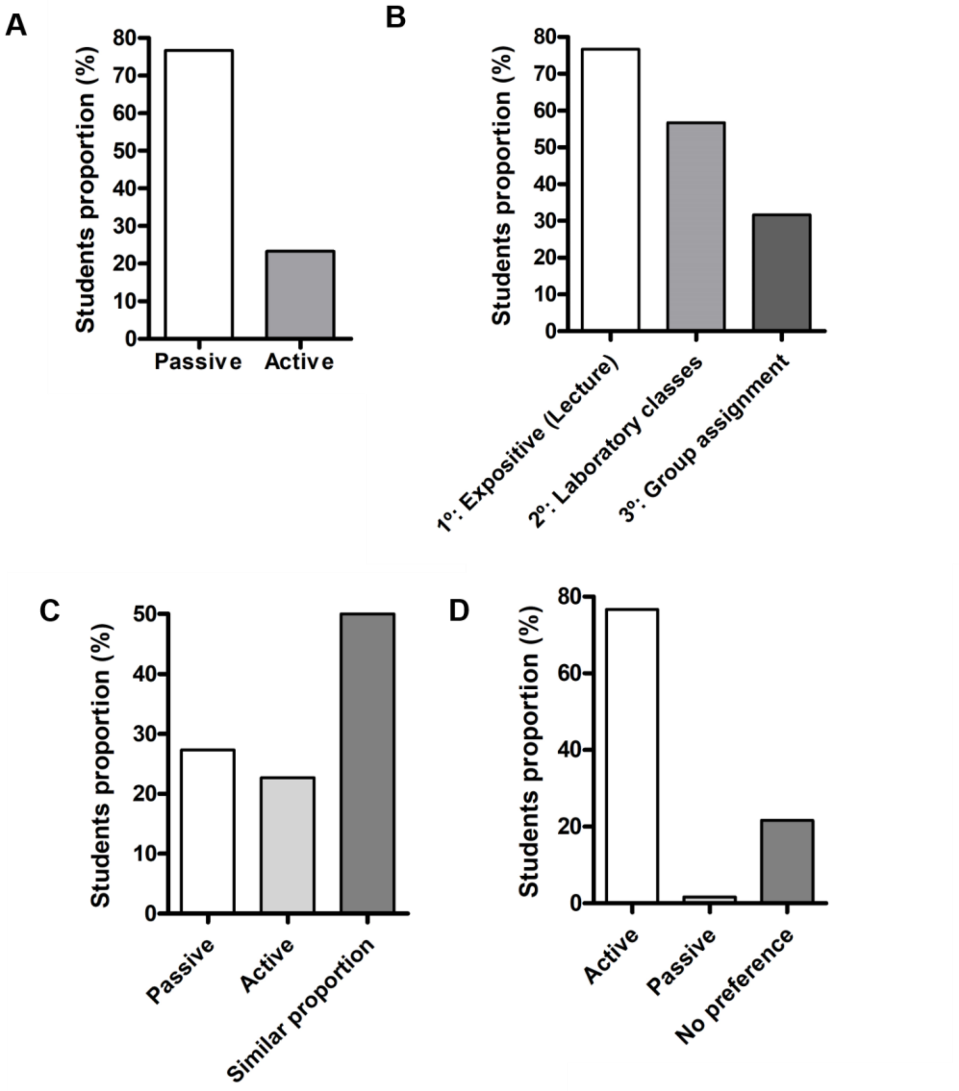


**Suppl. Fig. 4. Students' previous experience with teaching methodologies reveals a general profile of Brazilian students.** A. Teaching method (passive or active) most used during the participants' academic experience. B. Three main teaching approaches experienced by course participants. C. Predominance of the teaching approach during graduate classes (between passive or active). D. Evaluation of students regarding the effectiveness of active teaching methodologies in their learning.

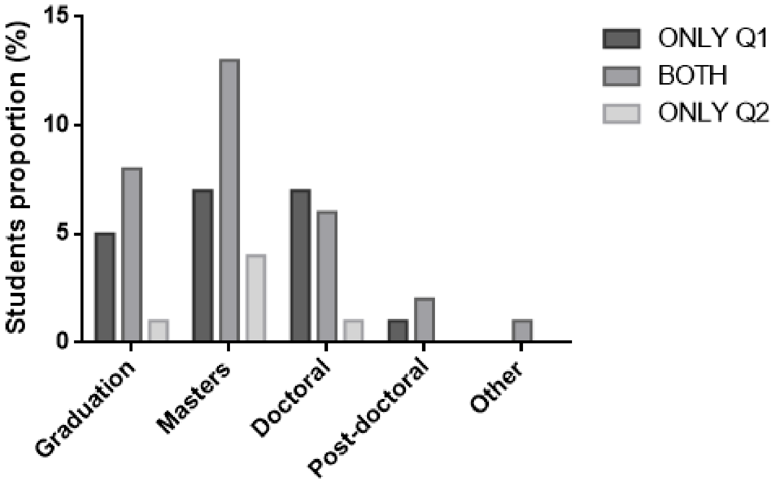


**Suppl. Fig. 5. Representation of participants engagement in questionnaires**. To analyze public diversity, students were separated in 3 subgroups according to the engagement in the questionnaires: subgroup “BOTH” (n=20). Students that participate only in one questionnaire were labelled as “ONLY Q1” (n=30) for the first questionnaire participation and “ONLY Q2” (n=6) for the second questionnaire participation.

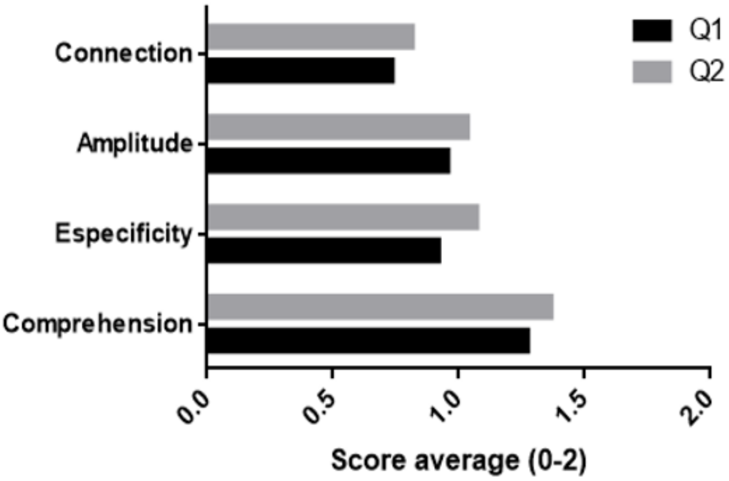


**Suppl. Fig. 6.** Students’ scores average at the four criteria used for open-ended questions evaluation;

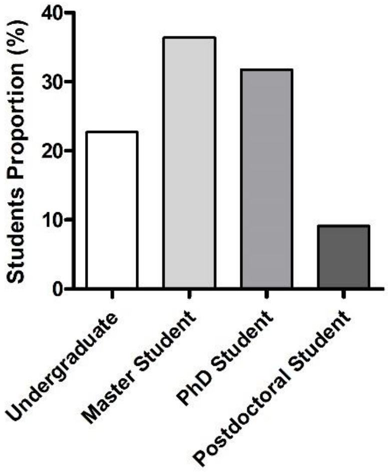


**Suppl. Fig. 7.** Academic background of students who submitted IRP.

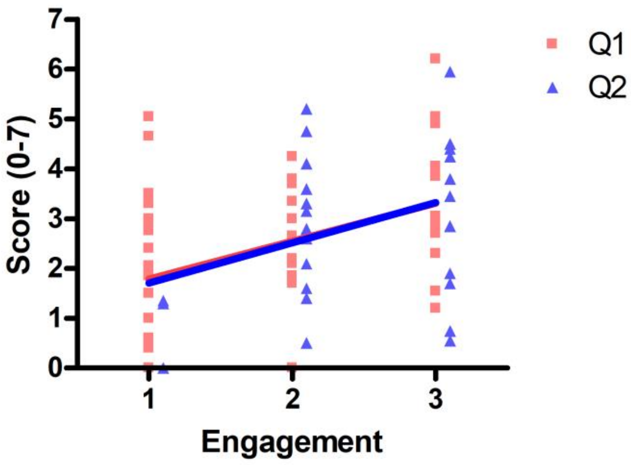


**Suppl. Fig. 8.** Correlation between level of engagement in course activities (Q1, Q2 and IRP. 1, 2 or 3) and performance assessed by the scores of the open-ended questions of Q1 (red) or Q2 (blue). The linear regression is shown by the slope. Correlation parameters are P value: Q1 = 0,081 and Q2 = 0,0046; and Person r: Q1 = 0,3987 and Q2 = 0,3443.

| | Q1 | Q2 | Improvement (%) |
|---|---|---|---|
| **Multiple-choice questions (0-1)** | 0.74 | 0.84 | 13,51% |
| **Open-ended questions (0-7)** | 2.36 | 2.73 | 15,67% |
| **General score (%)** | 59.96 | 61.74 | 2,96% |

**Suppl. Table 1 - Students' average scores improvement in Q2**. On average, students presented an improvement of 13,51% in objective and 15,67% in open-ended questions.

**Suppl. Video 1.** Moments of the course showing the dynamics of the course and the active methodology tools used. The video is available at: https://www.youtube.com/watch?v=jVM9AwJHJr8
